# Inositol hexakisphosphate is a critical regulator of Integrator assembly and function

**DOI:** 10.1101/2021.09.14.460315

**Authors:** Min-Han Lin, Madeline K. Jensen, Nathan D. Elrod, Kai-Lieh Huang, Eric J. Wagner, Liang Tong

**Affiliations:** Department of Biological Sciences, Columbia University, New York, NY 10027, USA; Department of Biochemistry and Molecular Biology, The University of Texas Medical Branch, Galveston, TX 77550, USA

## Abstract

Integrator has critical roles in noncoding RNA 3′-end processing as well as transcription attenuation of selected mRNAs. IntS11 is the endonuclease for RNA cleavage, as a part of the IntS4-IntS9-IntS11 complex (Integrator cleavage module, ICM). Our structure of the *Drosophila* ICM, determined by cryo-electron microscopy at 2.74 Å resolution, unexpectedly revealed the stable association of an inositol hexakisphosphate (IP_6_) molecule. The binding site is located in a highly electropositive pocket at an interface among all three subunits of ICM, 55 Å away from the IntS11 active site and generally conserved in other ICMs. IP_6_ binding is also confirmed in human ICM. Mutations of residues in this binding site or disruption of IP_6_ biosynthesis significantly reduced Integrator assembly and activity in snRNA 3′-end processing. Our structural and functional studies reveal that Integrator is subject to intricate cellular control and IP_6_ is a critical regulator of Integrator assembly and function in *Drosophila*, humans, and likely other organisms.

---

Integrator is a 14-subunit complex associated with RNA polymerase II (Pol II) that is crucial for 3′-end processing of snRNAs ^1^ and other noncoding RNAs ^2-5^, as well as transcription attenuation through cleavage of a subset of nascent mRNAs ^6-10^. The broad importance of Integrator function to gene regulation is evidenced by the wide range of human disease states attributed to dysfunction of its subunits ^11,12^. The Integrator subunits, named IntS1 through IntS14, can be purified as a complex but likely form several sub-modules. For example, our earlier studies have shown that IntS4-IntS9-IntS11 form the Integrator cleavage module (ICM) ^13,14^. IntS9 and IntS11 are paralogs of CPSF100 and CPSF73 in the canonical and U7 replication-dependent histone pre-mRNA 3′-end processing machineries, and CPSF73 catalyzes the cleavage reaction in both machineries ^15-17^. The subunits IntS10-IntS13-IntS14 form a nucleic acid binding module ^18,19^, and the IntS5-IntS8 complex ^19,20^ is critical for recruiting protein phosphatase 2A (PP2A) ^20-22^.

The structures of several Integrator components have been reported over the years, including the C-terminal domain (CTD2) complex of human IntS9-IntS11 ^23^, the N- and C-terminal domains of human IntS3 ^24,25^, human IntS13-IntS14 complex ^18^, and the human ICM ^19^. In addition, the structure of human Integrator in complex with PP2A was reported recently ^21^, in an inactive state. These structures reveal how Integrator is organized overall, but the mechanism of how RNA cleavage occurs is still unknown, and insight is lacking as to how Integrator activity, especially that of the endonuclease IntS11 ^1^, is regulated within this machinery and by other factors.

We co-expressed and purified the *Drosophila* ICM using baculovirus-infected insect cells (Figs. 1a,b) and determined its structure at 2.74 Å resolution by cryo-EM (Extended Data Figs. 1, 2a-c, Extended Data Table 1). Most of the residues of the three proteins in the complex could be located (Fig. 1a), with good side chain density. The overall structure of *Drosophila* ICM (Figs. 1c,d, Extended Data Fig. 2d) is generally similar to that of human ICM ^19,21^ (Extended Data Figs. 3a,b). IntS11 is in a closed, inactive state in our structure of *Drosophila* ICM, as well as those reported recently of human ICM ^19,21^. The structure shows that the C-terminal segments of IntS9 and IntS11 contain two separate domains, CTD1 and CTD2 (Figs. 1a,c), similar to their paralogs CPSF100 and CPSF73 ^17^. The two CTD2 domains have weak EM density (Extended Data Figs. 1, 2a-c), and their atomic models were guided by the structure of the human IntS9-IntS11 CTD2 complex ^23^. The CTDs have extensive interactions with each other, which should facilitate the association of IntS9 and IntS11. The metallo-β-lactamase and β-CASP domains of IntS9 and IntS11 form a pseudo-dimer in this structure (Fig. 1c), remarkably similar to the pseudo-dimer for the equivalent domains of CPSF100 and CPSF73 in the active U7 machinery (Extended Data Fig. 2e) ^17^. The N-terminal domain (NTD) of IntS4 contacts the metallo-β-lactamase domain of IntS9 and the back face of IntS11 metallo-β-lactamase and β-CASP domains (Fig. 1c), which may promote the formation of this pseudo-dimer.

**Figure 1.**
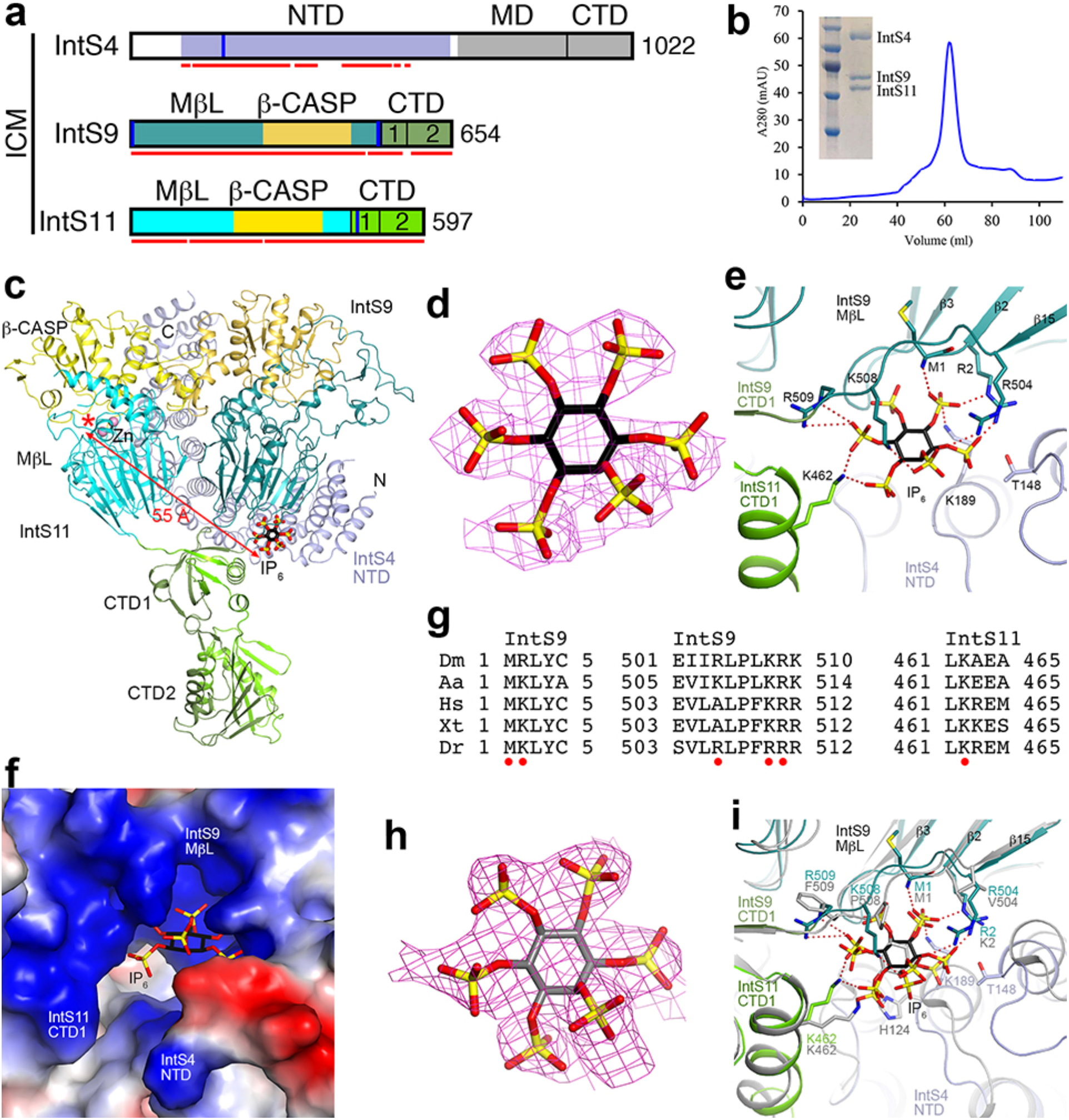
An IP_6_ binding site in the IntS4-IntS9-IntS11 complex (ICM). **(a)**. Domain organizations of *Drosophila* IntS4, IntS9 and IntS11. The domains are named and given different colors. The domains of IntS9 are shown in slightly darker colors compared to IntS11. Residues observed in the structure of the IntS4-IntS9-IntS11 complex are indicated with the red lines. The vertical bars in blue represent positively charged residues in the IP_6_ binding site. ICM: Integrator cleavage module; NTD: N-terminal domain; MD: middle domain; CTD: C-terminal domain. **(b)**. Gel filtration profile of *Drosophila* IntS4-IntS9-IntS11 complex. Inset: SDS gel of the purified complex. **(c)**. The overall structure of the *Drosophila* IntS4-IntS9-IntS11 complex. The domains are colored as in panel a and labeled. The IP_6_ molecule is shown as stick models (black for carbon atoms). **(d)**. Cryo-EM density for IP_6_. **(e)**. Detailed interactions between IP_6_ and the ICM. Ionic interactions between the phosphates of IP_6_ and ICM are indicated in dashed lines (red). **(f)**. Electrostatic surface of ICM near the binding site for IP_6_. **(g)**. Alignment of selected sequences for IntS9 and IntS11 residues in the IP_6_ binding site. Residues that have ionic interactions with the phosphate groups on IP_6_ are indicated with the red dots. Dm: *D. melanogaster*; Aa: *Aedes aegypti*; Hs: *H. sapiens*; Xt: *Xenopus tropicalis*; Dr: *Danio rerio*. **(h)**. Cryo-EM density for IP_6_ in human ICM (EMDB 12159) ^19^. An IP_6_ molecule was built into the density and subjected to real-space refinement. **(i)**. Overlay of the structures of *Drosophila* ICM (in color) and human ICM (gray) near the IP_6_ binding site. The human ICM structure likely has a registering shift of 2 residues for residues 504-509 of IntS9. Arg504, Lys508 and Arg509 of *Drosophila* IntS9 would actually be equivalent to Ala506, Lys510 and Arg511 instead of Val504, Pro508 and Phe509 of human IntS9, which would also be consistent with the sequence conservation in this region (panel g). The structure figures were produced with PyMOL (www.pymol.org) unless otherwise indicated.

**Figure 2.**
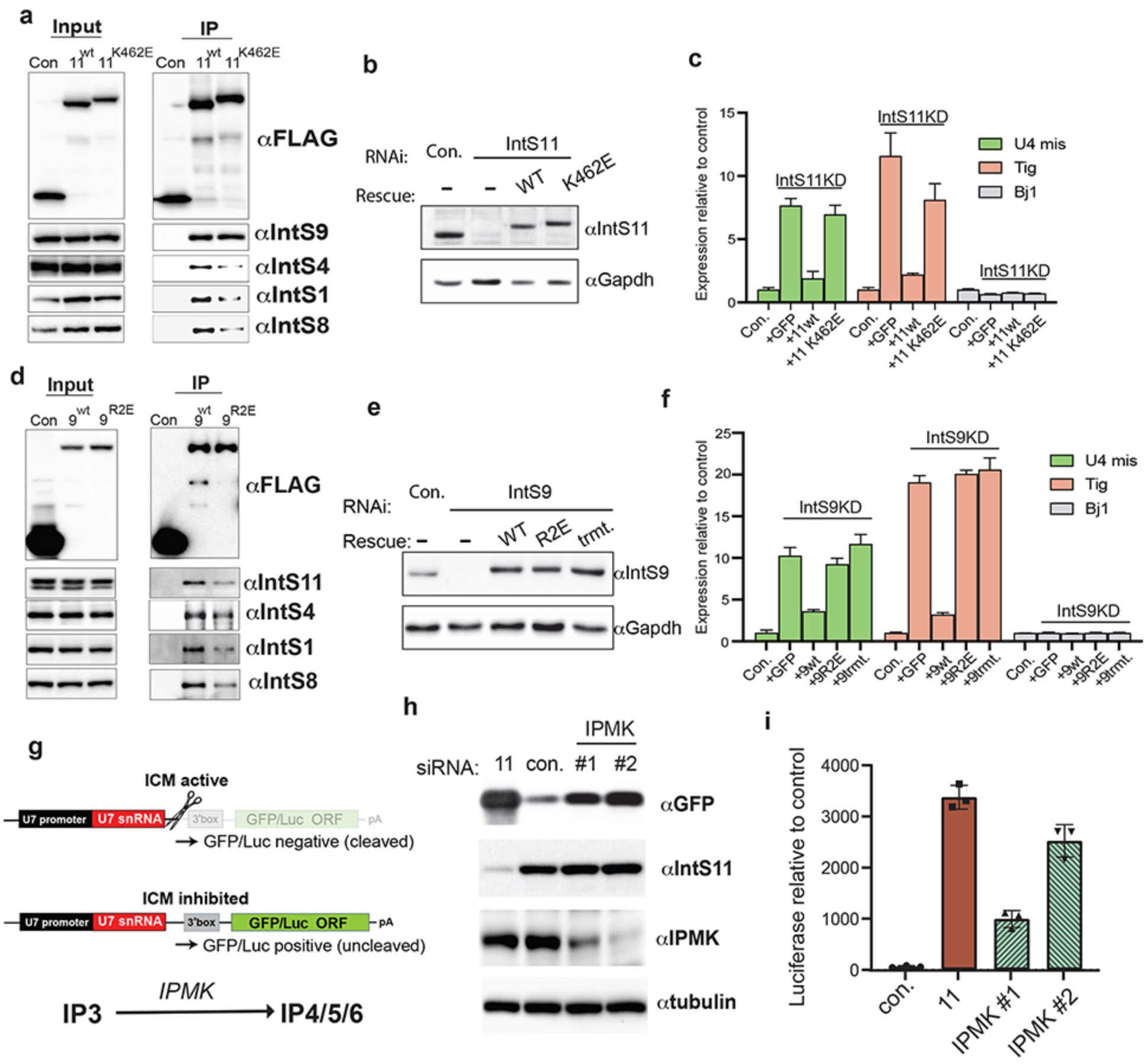
IP_6_ is required for Integrator Cleavage Module assembly and function. **(a)**. Western blot analysis of input nuclear extracts (left) and IP (right) from DL1 cells stably expressing FLAG-tagged proteins as indicated. IP conducted using anti-FLAG affinity resin was normalized to FLAG signals in each IP. **(b)**. Western blot analysis of DL1 cell lysates derived from cells treated with either control or IntS11 dsRNA. Rescues were conducted in cells depleted of IntS11 through inducible expression of RNAi-resistant cDNA. **(c)**. RT-qPCR analysis of selected genes identified to be upregulated upon IntS11 depletion using RNA-seq. Results are shown from biologically independent replicates, depicting averages and standard deviations (mean±SD, n=3). **(d)**. Western blot analysis of input nuclear extracts (left) and IP (right) from DL1 cells stably expressing FLAG-tagged proteins as indicated. **(e)**. Western blot analysis of DL1 cell lysates derived from cells treated with either control or IntS9 dsRNA. Rescues were conducted in cells depleted of IntS9 through inducible expression of RNAi-resistant cDNA. **(f)**. RT-qPCR analysis of selected genes identified to be upregulated upon IntS11 depletion using RNA-seq. Results are shown from biologically independent replicates, depicting averages and standard deviations (mean±SD, n=3). **(g)**. Schematic of the U7snRNA reporter systems, where the U7 gene is cloned upstream of GFP or luciferase. Top: in WT cells, the ICM drives RNA Pol II termination and prevents GFP/luc expression. Bottom: upon loss of ICM function through IPMK depletion, RNA Pol II productively elongates through the GFP/luc ORF, yielding expression. **(h)**. Western blot analysis of 293T cell lysates treated with siRNA as labeled as well as the U7-GFP reporter. The levels of IntS11 and IPMK depletion are shown as well as the level of GFP expression from the reporter. **(i)**. Similar experiment as in panel h except the U7-luc reporter was used and light units were measured in each case.

The structure of *Drosophila* ICM unexpectedly revealed the presence of an inositol hexakisphosphate (IP_6_) molecule (Fig. 1c), with good quality EM density (Fig. 1d). The compound was bound to the ICM during expression in insect cells and remained associated through two steps of column purification, suggesting that it has high affinity for ICM. IP_6_ is located at an interface among all three subunits of the ICM, formed by the N-terminus and the linker to CTD1 of IntS9, the CTD1 of IntS11, and the first few helical repeats of IntS4 NTD (Fig. 1e). IP_6_ has ionic interactions with all three subunits, including Lys189 of IntS4, the N-terminal ammonium ion, Arg2, Arg504, Lys508 and Arg509 of IntS9, and Lys462 of IntS11 (Fig. 1e). These residues create a large, highly positively charged pocket (Fig. 1f), suggesting that the binding of IP_6_ here may help stabilize the conformation of ICM.

Residues in the IP_6_ binding site are generally conserved among IntS9 and IntS11 homologs (Fig. 1g). While Lys189 is a glutamine in vertebrate IntS4, IP_6_ should maintain the favorable interactions with the dipoles of the helices in the IntS4 NTD. In the structure of the human ICM ^19^, EM density highly consistent with IP_6_ is present near the N-terminus of IntS9 as well (Fig. 1h), although IP_6_ was not included in that atomic model. The detailed binding mode of IP_6_ in human ICM has only slight differences compared to that in *Drosophila* ICM (Fig. 1i). In the structure of human Integrator-PP2A complex (EMDB 30473) ^21^, some EM density is present near the N-terminus of IntS9, although the quality of the density here is poor. In fact, a few N-terminal residues of IntS9 were built into this density. Overall, the structural observations suggest that this pocket is likely to bind IP_6_ or another negatively charged compound(s) and play a role in the function of ICM in general.

To characterize the importance of IP_6_ in Integrator function, we created mutations in the IP_6_ binding site that changed the positively charged residues to negatively charged ones and assessed their impact on assembly and function of Integrator. We created DL1 nuclear extracts from cell lines stably expressing FLAG-IntS11-WT or FLAG-IntS11-K462E and then purified associated complexes using anti-FLAG affinity. We observed that the K462E mutation greatly reduced association with IntS4, IntS1, and IntS8 but did not significantly alter the binding to IntS9 (Fig. 2a). To determine the functional significance of IP_6_ binding to Integrator, we devised a method to induce expression of IntS11-WT or IntS11-K462E in cells where endogenous IntS11 has been depleted using dsRNA targeting its 5′ and 3′ UTRs. We observed that treatment of DL1 cells with IntS11 dsRNA resulted in effective depletion of endogenous IntS11, whereas induction of IntS11 transgenes allowed expression of near endogenous levels of IntS11-WT or IntS11-K462E proteins (Fig 2b). We then analyzed the impact of IntS11 depletion on U4snRNA processing, transcriptional attenuation of a previously validated mRNA-encoding gene called tigrin (tig) ^6^, and a gene found not to be regulated by Integrator (Bj1). As expected, depletion of IntS11 resulted in a significant increase in U4snRNA misprocessing as well as tig transcription but had no effect on Bj1 (Fig. 2c). Importantly, these phenotypes were restored upon re-expression of IntS11-WT but were found not to be rescued by the IntS11-K462E mutant (Fig. 2c).

To further probe the importance of IP_6_ to Integrator assembly and function, we created nuclear extracts from DL1 cells expressing either FLAG-IntS9-WT or FLAG-IntS9-R2E, which is predicted to disrupt IP_6_ interaction. We purified associated complexes using anti-FLAG affinity resin and found that the R2E mutation greatly reduced association with other Integrator subunits (Fig. 2d). To address the functional importance of the IntS9 residues involved in IP_6_ binding, we repeated a similar RNAi rescue experiment as was done for IntS11. We found that treatment of DL1 cells with IntS9 dsRNA resulted in effective depletion of endogenous IntS9, whereas induction of RNAi-resistant IntS9 transgenes allowed expression of near endogenous levels of IntS9-WT, IntS9-R2E, or an IntS9 containing mutations at all three basic residues located at 504, 508 or 509 (IntS9-trmt) (Fig. 2e). Depletion of IntS9 led to similar gene expression changes to that of IntS11 depletion and that re-expression of IntS9-WT could reverse these changes (Fig. 2f). Importantly, expression of the IntS9-R2E or IntS9-trmt did not rescue the depletion phenotype, indicating that these residues are required for Integrator function (Fig. 2f).

Finally, we transfected cells with two different siRNAs targeting inositol polyphosphate multikinase (IPMK), which is required for IP_6_ biosynthesis ^26,27^. We assessed Integrator function using two reporters where the U7snRNA promoter, gene body, and 3′ cleavage site are placed upstream of either GFP or luciferase with the rationale that loss of Integrator function leads to transcriptional readthrough and expression of the downstream open reading frames (Fig. 2g). We observed that depletion of IPMK resulted in increased expression of GFP and luciferase relative to control (Figs. 2h,i). Notably, the siRNA capable of more significant IPMK depletion (#2 in Figs. 2h,i) produced much higher GFP/luciferase expression, in fact reaching a level similar to that observed after depletion of IntS11.

The binding site for IP_6_ in ICM is located far (55 Å) from the active site of IntS11 and is on the opposite face from the opening for the canyon in IntS11 for binding RNA (Fig. 1c). Therefore, this binding site is unlikely to have a direct effect on the catalysis by IntS11. Mutation of Lys462 also affected human Integrator function, and it was suggested this binding site could interact with a part of the snRNA substrate ^19^, although our functional studies have demonstrated a critical role for IP_6_ in Integrator activity. Therefore, it is more likely that this is a site for binding allosteric modulators (such as IP_6_) of ICM, revealing a previously unknown mechanism of regulating Integrator function. Our observations also expand the repertoire of IP_6_ as a signaling molecule impacting RNA processing ^28-32^ as well as other processes in the cell ^33,34^.

## Methods

### Protein expression and purification

*Drosophila* IntS4, IntS9, and IntS11 were co-expressed in insect cells. IntS9 and IntS11 were cloned into the pFL acceptor vector. N-terminal 6xHis-tagged IntS4 was cloned into the pSPL donor vector. These two vectors were fused by Cre recombinase. High5 cells (2 × 10^6^ cells·ml^−1^) were infected with 16 ml of IntS4-IntS9-IntS11 P2 virus and harvested after 48 h.

For purification, the cell pellet was resuspended and lysed by sonication in 100 ml of buffer containing 20 mM Tris (pH 8.0), 250 mM NaCl, 2 mM βME, 5% (v/v) glycerol, and one tablet of protease inhibitor mixture (Sigma). The cell lysate was then centrifuged at 13,000 rpm for 40 min at 4 °C. The protein complex was purified from the supernatant via nickel affinity chromatography. The protein complex was further purified by a Hiload 16/60 Superdex 200 column. The IntS4-IntS9-IntS11 complex was concentrated to 2 mg·ml^−1^ in a buffer containing 20 mM Tris (pH 8.0), 300 mM NaCl, and 2 mM DTT, and stored at –80 °C.

### EM specimen preparation and data collection

All specimens for cryo-EM were frozen with an EM GP2 plunge freezer (Leica) set at 20 °C and 99% humidity. Cryo-EM imaging was performed in the Simons Electron Microscopy Center at the New York Structural Biology Center using Leginon ^35^.

For the IntS4-IntS9-IntS11 complex, a 3.5 μL aliquot at 0.1 mg·ml^−1^ was applied to one side of a Quantifoil 400 mesh 1.2/1.3 gold grid with graphene oxide support film (Quantifoil). After 30 s, the grid was blotted for 1.5 s on the other side and plunged into liquid ethane. 3,083 image stacks were collected on a Titan Krios electron microscope at New York Structural Biology Center, equipped with a K3 direct electron detector (Gatan) at 300 kV with a total dose of 51 e^−^ Å^−2^ subdivided into 40 frames in 2 s exposure using Leginon. The images were recorded at a nominal magnification of 81,000× and a calibrated pixel size of 1.083 Å, with a defocus range from –1 to −2.5 μm.

### Image processing

Image stacks were motion-corrected and dose-weighted using RELION 3.1 ^36^. The patch CTF parameters were determined with cryoSPARC ^37^. First, 2,920,144 particles were auto-picked and subjected to 2D classification in cryoSPARC. 1,149,821 particles in classes with recognizable features by visual inspection were used to generate eight 3D initial models by *ab initio* reconstruction. After one round of heterogeneous refinement, 620,438 particles were imported to RELION for CTF refinement and Bayesian polishing, yielding a map at 2.74 Å resolution.

### Model building

Atomic models for IntS11, IntS9 and IntS4 were built manually into the cryo-EM density with Coot ^38^. Homology models for *Drosophila* IntS9 and IntS11 were generated with I-TASSER ^39^, based on the structures of human CPSF100 and CPSF73 ^40^. The atomic models were improved by real-space refinement with the program PHENIX ^41^.

The model of IP_6_ in the *Drosophila* ICM was placed in the human ICM cryo-EM density (EMDB 12159) ^19^ and manually adjusted to fit the density. Real-space refinement was then used to optimize the fitting of IP_6_ to the EM density.

### Plasmid construction and stable cell lines generation

For mutation analysis of *Drosophila* IntS9 and IntS11, site-directed PCR mutagenesis was used to create the IntS9-R2E, IntS9-trmt, and IntS11-K462E mutations. Wild-types and the mutants of dIntS9 and IntS11 were subsequently cloned into the pMT-3xFLAG-puro vector ^6^ to inducibly express in DL1 cells. All plasmids were sequenced to confirm identity. To generate cells stably expressing the FLAG-IntS9-WT, FLAG-IntS9-R2E, FLAG-IntS9-trmt, FLAG-IntS11-WT, FLAG-IntS11-K462E, and eGFP control transgenes, 2×10^6^ cells were first plated in regular maintenance media in a 6-well dish overnight. 2 μg of expressing plasmids were transfected using Fugene HD (Promega, #E2311). After 24 hours, 2.5 μg/mL puromycin was added to the media to select and maintain the cell population.

### Nuclear extract preparation

Five 150 mm dishes of each condition of confluent cells (pretreated with 500 μM CuSO_4_ for 24 hours) were collected and washed in cold PBS before being resuspended in ten times volumes of the cell pellet of Buffer A (10 mM Tris pH 8, 1.5 mM MgCl_2_, 10 mM KCl, 0.5 mM DTT, and 0.2 mM PMSF). Resuspended cells were allowed to swell during a 15-minute rotation at 4 °C. After pelleting down at 1,000 g for 10 minutes, two times volumes of the original cell pellet of Buffer A were added and cells were homogenized with a dounce pestle B for 40 strokes on ice. Nuclear and cytosolic fractions were then separated by centrifuging at 800 g for 10 minutes. To attain a nuclear fraction, the pellet was washed once with Buffer A before being resuspended in two times volumes of the original cell pellet of Buffer C (20 mM Tris pH 8, 420 mM NaCl, 1.5 mM MgCl_2_, 25% (v/v) glycerol, 0.2 mM EDTA, 0.5 mM PMSF, and 0.5 mM DTT). The samples were then homogenized with a dounce pestle B for 20 strokes on ice and rotated for 30 minutes at 4 °C before centrifuging at 15,000 g for 30 minutes at 4 °C. Finally, supernatants were collected and subjected to dialysis in Buffer D (20 mM HEPES, 100 mM KCl, 0.2 mM EDTA, 0.5 mM DTT, and 20% (v/v) glycerol) overnight at 4 °C against 3.5 kDa MWCO membrane (Spectrum Laboratories, #132720). Prior to any downstream applications, nuclear extracts were centrifuged again at 15,000 g for 3 minutes at 4 °C to remove any precipitate.

### Western blotting and anti-FLAG affinity purification

To check protein expression, cells were lysed directly in wells in 2X SDS sample buffer (120 mM Tris pH 6.8, 4% SDS, 200 mM DTT, 20% (v/v) glycerol, and 0.02% bromophenol blue). Lysates were incubated at room temperature with periodic swirling prior to a 10-minute boiling at 95 °C and a short sonication. Denatured protein samples were then resolved in a 10% SDS-PAGE and transferred to a PVDF membrane (Bio-Rad, #1620177). Blots were probed by custom-designed *Drosophila* antibodies as previously described ^6^ diluted in PBS-0.1% Tween supplemented with 5% nonfat milk. To detect proteins from 293T lysate, anti-hInts11 (Bethyl, #A301-274A), anti-hIMPK (Thermo, #PA5-21629), anti-GFP (Clontech, #632381), anti-alpha Tubulin (abcam, #ab15246), and anti-GAPDH (Thermo, #MA5-15738) were used at the dilution suggested by the manufacturer.

To purify FLAG-tagged Integrator complexes, 1 mg of nuclear extract was mixed with 40 μL anti-Flag M2 affinity agarose slurry (Sigma, #A2220) equilibrated in binding buffer (20 mM HEPES pH 7.4, 100 mM KCl, 10% (v/v) glycerol, 0.1% NP-40) and rotated for 2 hours at 4 °C. Following the two-hour incubation/rotation, five sequential washes were carried out in binding buffer with a 10-minute rotation at 4 °C followed by a 500 g centrifugation at 4 °C. After the final wash, the binding buffer supernatant was removed using a pipette and the protein complexes were eluted from the anti-FLAG resin by adding 40 μL of 2X sample buffer and boiled at 95 °C for five minutes. For input samples, nuclear extracts were mixed with 5X loading buffer and boiled, and 1/10 volume of the immunoprecipitation reaction was loaded on SDS-PAGE.

### RNA Interference

Double-stranded RNAs targeting the 5′ and 3′ UTRs of *Drosophila* IntS11 and an RNAi resistant region of the IntS9 constructs were generated by *in vitro* transcription of PCR templates containing the T7 promoter sequence on both ends using MEGAscript kit (Thermo, #AMB13345). For RNA interference experiments, 1.5×10^6^/ml of DL1 cells were washed into serum free media and seeded into a 6-well plate along with 10 μg of dsRNA. After a 1-hour incubation, 2 mls of complete growth medium was added followed by 60 hours of incubation before harvest. To perform rescue experiments while knocking down, cells were also treated with 100 mM CuSO_4_ throughout the 60 hours incubation period to induce expression of the RNAi-resistant FLAG-IntS9-WT, FLAG-IntS9-R2E, FLAG-IntS9-trmt, FLAG-IntS11-WT, FLAG-IntS11-K462E transgenes.

IntS11 (Sigma, #SASI_Hs01_00032429), IPMK (Sigma, #SASI_Hs01_00047017 and #SASI_Hs01_00047015), and control (Sigma, #SIC002) siRNAs (2 μL each of 20 mM stock) were incubated in 50 μL of pre-warmed (room temperature) Opti-MEM I reduced serum medium (GIBCO) for 5 minutes at room temperature. Similarly, RNAiMax (2 μL per well) was incubated in 50 μL pre-warmed Opti-MEM I reduced serum medium for 5 minutes. The siRNA and RNAiMAX dilutions were mixed and incubated for 20 minutes at room temperature. 2×10^5^ 293T cells were seeded into a 24-well plate and the prepared transfection mixes of 40 pmols siRNAs were added to each well. Cells were transfected with 60 pmols of siRNA second time after 24 hours of incubation. The cells were expanded into 12-well plate after total 48 hours, and harvested at total 72 hours of incubation under standard mammalian cell culture conditions.

### Reporter cell lines establishment and luciferase measurement

To construct a reporter plasmid, pAAVS1-TLR targeting vector (Addgene, #64215) was cut with CalI and PspXI to substitute with U11/U7 small nuclear RNA (500 bp upstream of transcription start site, coding region, and 50 bp downstream of coding region) followed by Renilla luciferase/GFP coding region and SV40 poly(A) signal. The reporter plasmid was co-transfected with the gRNA, cloned into pU6-(BbsI) CBh-Cas9-T2A-mCherry (Addgene, #64324), targeting AAVS1 locus into 293T. Briefly, equal amount of plasmids were transfected with lipofetamine 2000 for 24 hours before selecting with 800 ng/ml puromycin for 2 days. Cells were then grown in regular medium without puromycin for a week before clonal selection.

Renilla Luciferase Assay System (Promega, #E2810) was used to assess luciferase activity in the clonal reporter cell lines. The line with highest luciferase activity after hIntS11 knock down was selected. To test the role of IPMK in Integrator’s function, the reporter cell lines were seeded in 24-well plate and knocked down twice with siRNAs as described. Luciferase activity was measured according to manufacturer’s protocol. Background luciferase activity of each sample was calculated by interpolation and subtracted. Briefly, scatterplot and trendline were plotted by protein quantity versus luciferase activity obtained from the same amount of reporter cell line lysate, and increased amount of the lysate were measured. The bars represent the average of triplicate biological repeats.

### RT-qPCR quantification and analysis

Data were analyzed using the DDCt method with Rps17 as the reference gene and LacZ dsRNA-treated cells as the control, as described previously^42^.

## Acknowledgments

We thank Oliver Clarke for helpful discussions, the staff at the Columbia University Cryo-Electron Microscopy Center for help with screening EM grids, Huihui Kuang and the staff at the New York Structural Biology Center for help with cryo-EM data collection. This research is supported by the Cancer Prevention Research Institute of Texas (CPRIT) grant RP170593 (to K.-L.H.), a Kempner Predoctoral Fellowship (to N.D.E.), NIH grants R01GM134539 (to E.J.W) and R35GM118093 (to L.T.).

## Author Contributions

N.D.E., M.K.J., and K-L.H. conducted biochemical purifications, immune-precipitations and western blotting; M-H.L. and L.T. conducted all structural analyses; N.D.E. and M.K.J. generated all constructs used in DL1/293T cells; W.K.R. performed all mass spectrometry; E.J.W. and L.T. conceived the project and wrote the manuscript, with comments from all authors.

## Data availability

The structure of *Drosophila* ICM-IP_6_ complex will be deposited at the PDB for release upon publication.

