## Supplementary Information for "Inositol hexakisphosphate is a critical regulator of Integrator assembly and function"

**Extended Data Table 1**
**Cryo-EM data collection, refinement and validation statistics**

| | IntS4-IntS9-IntS11 complex |
|---|---|
| **Data collection and processing** | |
| Magnification | 81,000 |
| Voltage (kV) | 300 |
| Electron exposure ($e^{-}$/Å$^2$) | 51 |
| Defocus range (μm) | -1 to -2.5 |
| Pixel size (Å) | 1.083 |
| Symmetry imposed | C1 |
| Image stacks (no.) | 3,083 |
| Initial particles images (no.) | 2,920,144 |
| Final particle images (no.) | 620,438 |
| Map resolution (Å) | 2.74 |
| FSC threshold | 0.143 |
| Map sharpening B-factor (Å$^2$) | –81.8 |
| **Refinement** | |
| Number of protein residues | 1,621 |
| Number of metal ions | 2 |
| Number of atoms | 12,819 |
| R.m.s. deviations | |
| Bond lengths (Å) | 0.010 |
| Bond angles (°) | 1.093 |
| PDB validation | |
| Clash score | 6.34 |
| Poor rotamers (%) | 0.07 |
| Ramachandran plot | |
| Favored (%) | 90.11 |
| Allowed (%) | 9.89 |
| Disallowed (%) | 0.00 |

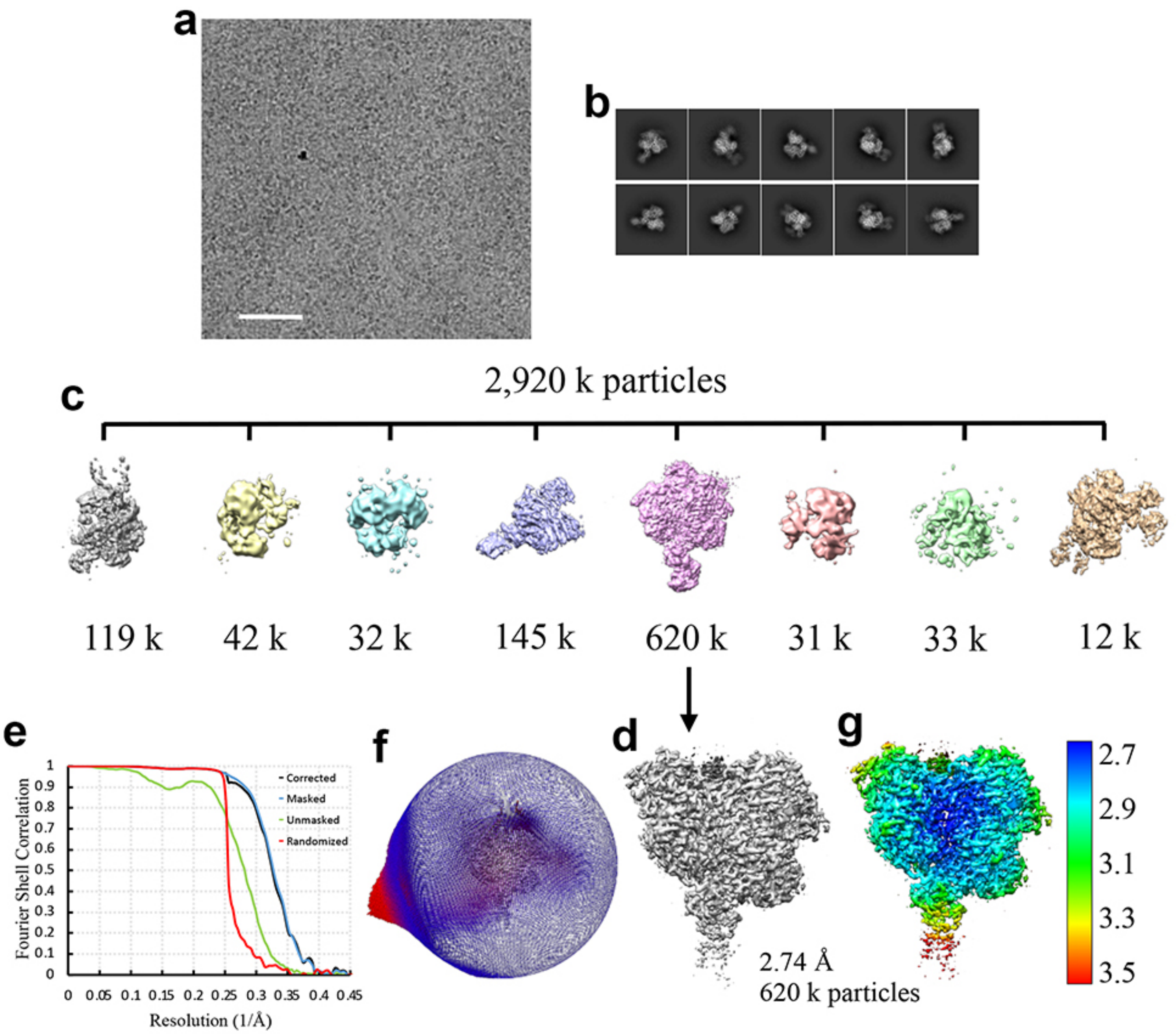

**Extended Data Fig. 1. Single-particle cryo-EM analysis of the *Drosophila* IntS4-IntS9-IntS11 complex. (a).** A region of a cryo-EM micrograph of the IntS4-IntS9-IntS11 complex. Scale bar: 50 nm. **(b).** Ten 2D class averages from the cryo-EM micrographs. **(c).** 3D classification (hetero refinement) of particles identified from 2D class averages. **(d).** Final cryo-EM map after Bayesian polishing and refinement in RELION. **(e).** Fourier shell correlation curves for the final model. **(f).** Orientations of particles used in the refinement for the final cryo-EM reconstruction. **(g).** Local resolution map for the final cryo-EM reconstruction.

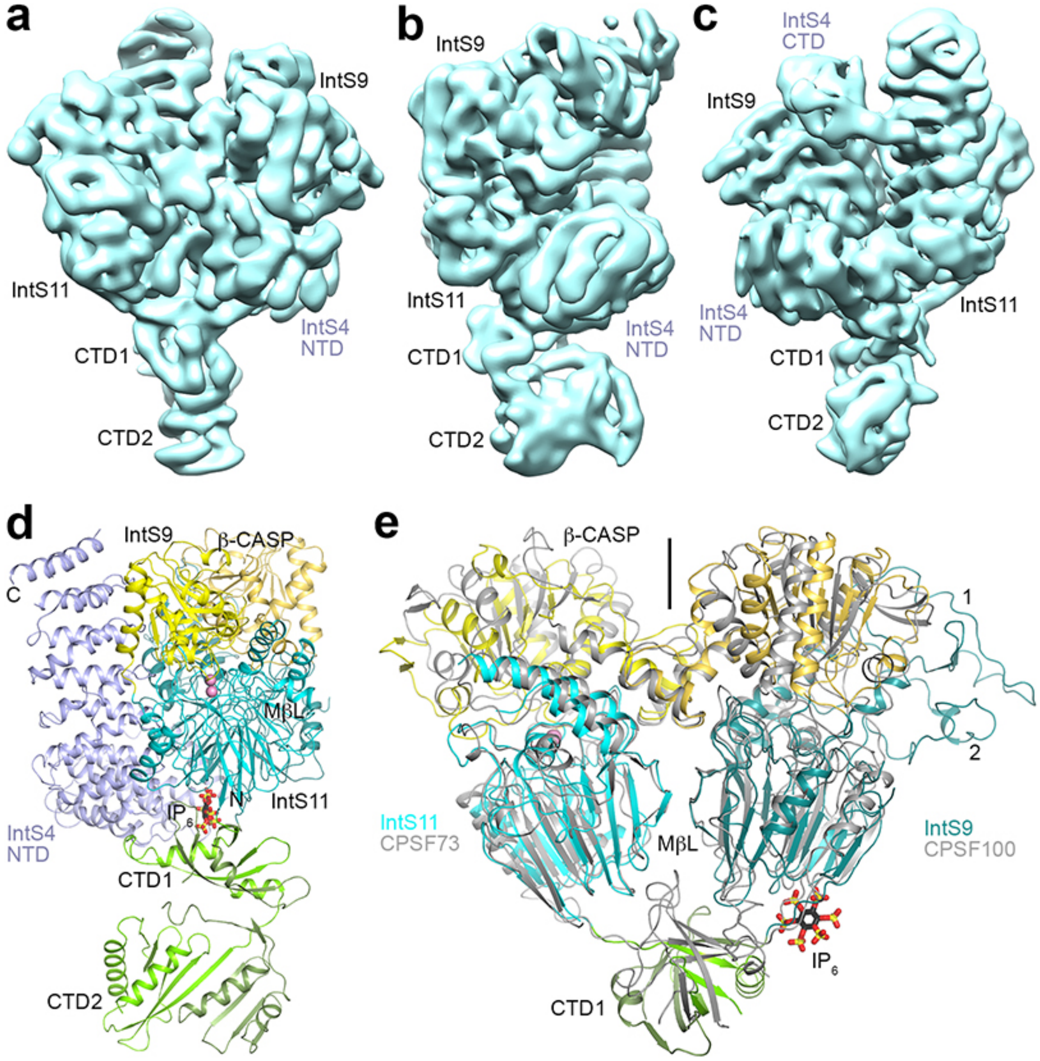

**Extended Data Fig. 2. The IntS9-IntS11 dimer is similar to that of the CPSF100-CPSF73 dimer**. **(a)**. Cryo-EM density of the IntS4-IntS9-IntS11 complex low-pass filtered to 7 Å resolution, and viewed after 90° **(b)** and 180° **(c)** rotation around the vertical axis. Weak EM density for CTD2 of IntS9 and IntS11 and residues beyond 570 of IntS4 (CTD) becomes more visible at this resolution, but the quality of the density was not sufficient to allow an atomic model to be built for IntS4 CTD. **(d)**. Another view of the structure of the IntS4-IntS9-IntS11 complex, related to that of Fig. 1c by a 90° rotation around the vertical axis. **(e)**. Overlay of the dimer of the metallo-β-lactamase and β-CASP domains of IntS9-IntS11 (in color) in the ICM with the dimer of the equivalent domains of human CPSF100-CPSF73 (gray) in the active histone pre-mRNA 3′-end processing machinery. The pseudo two-fold axis of this dimer is along the vertical direction (indicated with the black line). The CTD1s of the two structures assume very different positions, and the CTD2s are not shown. IntS9 metallo-β-lactamase domain contains two insertions, residues 34-83 (labeled 1) and 150-179 (2), that are positioned next to the β-CASP domain.

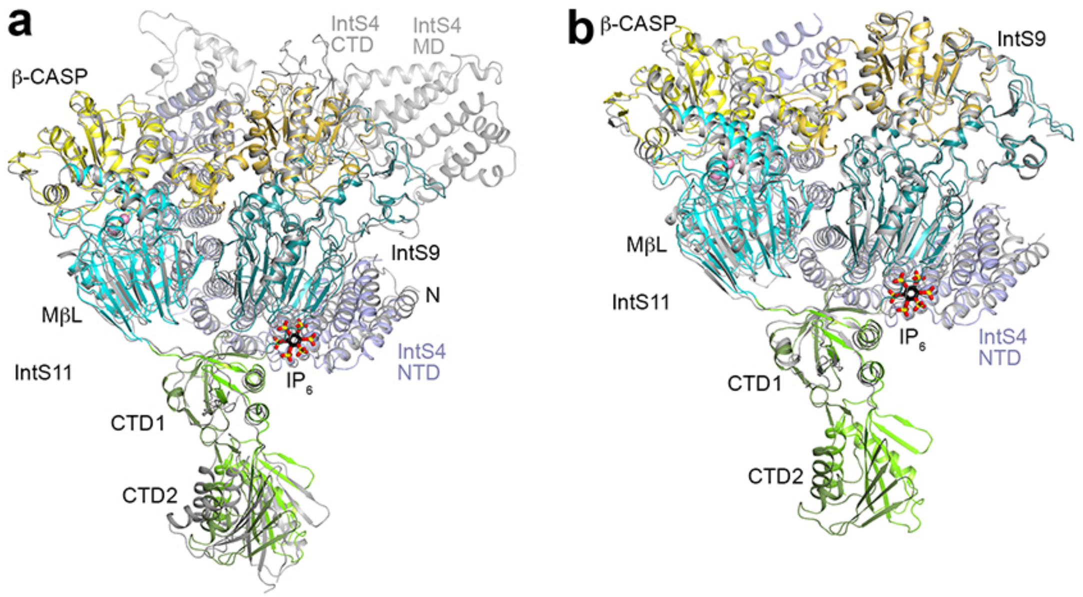

**Extended Data Fig. 3. Structural comparison between *Drosophila* and human ICM**. **(a).** Overlay of the overall structure of *Drosophila* ICM and human ICM in the structure of Integrator in complex with PP2A (PDB entry 7CUN). The positions of CTD2 of IntS9 and IntS11 are noticeably different, likely related to the flexibility in these domains. There are differences in the registering of residues in places, and many of the residues in human Integrator lack models for the side chain due to poor EM density. **(b)**. Overlay of the overall structure of *Drosophila* ICM and human ICM (PDB entry 7BFP). The CTD2 of IntS9 and IntS11 are not modeled in the human ICM. Differences in registering are also observed, for example see Fig. 1i.
